# Theta power and theta-gamma coupling support spatial memory retrieval

**DOI:** 10.1101/732735

**Authors:** Umesh Vivekananda, Daniel Bush, James A Bisby, Sallie Baxendale, Roman Rodionov, Beate Diehl, Fahmida A Chowdhury, Andrew W McEvoy, Anna Miserocchi, Matthew C Walker, Neil Burgess

## Abstract

Hippocampal theta oscillations have been implicated in spatial memory function in both rodents and humans. What is less clear is how hippocampal theta interacts with higher frequency oscillations during spatial memory function, and how this relates to subsequent behaviour. Here we asked ten human epilepsy patients undergoing intracranial EEG recording to perform a desk-top virtual reality spatial memory task, and found that increased theta power in two discrete bands (‘low’ 2-5Hz and ‘high’ 6-9Hz) during cued retrieval was associated with improved task performance. Similarly, increased coupling between ‘low’ theta phase and gamma amplitude during the same period was associated with improved task performance. These results support a role of theta oscillations and theta-gamma phase-amplitude coupling in human spatial memory function.

## Introduction

Oscillations in the local field potential (LFP) reflect synchronous neural activity and are a likely candidate to integrate functional brain regions across multiple spatiotemporal scales (Buzsáki & Schomburg, 2015; Fries, Nikolic, & Singer, 2007). In particular, oscillations within the hippocampal-entorhinal system have long been hypothesized to play a role in cognitive function. The theta rhythm has been well documented in the rodent and human hippocampal network during translational movement and memory function (Buzsáki & Moser, 2013; Düzel, Penny, & Burgess, 2010; O’Keefe & Nadel, 1978; Vanderwolf, 1969). Theta frequency in rodents is typically 6-12Hz, but in the human hippocampus, theta frequency appears to be lower and occupy discrete ‘low’ (2-5Hz) and ‘high’ (6-9Hz) bands (Bush et al., 2017; Lega, Jacobs, & Kahana, 2012; Watrous, Tandon, Conner, Pieters, & Ekstrom, 2013). The modulation of high-frequency activity by the phase of low-frequency oscillations such as theta, manifesting as phase amplitude coupling (PAC), may provide a mechanism for inter-areal communication and phase coding (Canolty et al., 2006). PAC has been well documented in both human and animal studies during spatial (Bieri, Bobbitt, & Colgin, 2014; Lisman & Jensen, 2013; Newman, Gillet, Climer, & Hasselmo, 2013; Tamura, Spellman, Rosen, Gogos, & Gordon, 2017; Tort et al., 2008), declarative (Axmacher et al., 2010; Fell et al., 2003; Lega, Burke, Jacobs, & Kahana, 2016; Tort, Komorowski, Manns, Kopell, & Eichenbaum, 2009), and sequence memory tasks (Heusser, Poeppel, Ezzyat, & Davachi, 2016). In particular, the modulation of low (30-50Hz) and high gamma (60-100Hz) power by theta phase has been described in both the rodent (Colgin, 2015; Colgin et al., 2009) and human brain (Alekseichuk, Turi, Amador de Lara, Antal, & Paulus, 2016; Lega et al., 2012). Here, we characterised the role of low and high theta oscillations, and their relationship with concurrent gamma power, in human intracranial EEG recordings during a self-paced spatial memory task. We found that low and high theta power and low and high gamma power were significantly increased during spatial memory retrieval, and that both increased theta power and increased PAC between low theta and gamma oscillations in the hippocampus during spatial memory retrieval correlated with task performance. These results support the hypothesis that theta-gamma PAC within the hippocampal formation contributes to successful spatial memory retrieval in humans.

## Methods

### iEEG Recordings

Thirteen patients with drug refractory epilepsy undergoing intracranial EEG monitoring for clinical purposes were asked to perform a spatial memory task. Single group comparison analysis based on the effect size observed in a previous translational movement study (see Bush et al., 2017) indicates that 10 patients would be required to identify differences in theta power at p < 0.05 with a power of 90%. Prior approval was granted by the NHS Research Ethics Committee, and informed consent was obtained from each subject. Post-implantation CT and/or MRI scans were used to visually inspect and identify electrode locations, confirming that 10 patients had hippocampal contacts. Patient demographics are listed in Supplemental Table 1. Depth EEG was recorded continuously at a sample rate of 1,024 Hz (patients 4 and 10) or 512 Hz (all other patients) using either a NicoletOne long-term monitoring system (Natus Medical, Inc.) (Patients 1–6) or Micromed SD long-term monitoring system (Micromed) (patients 7–10). Recordings made at a higher sampling rate were down-sampled to 512 Hz, to match those from the majority of patients, before any analyses were performed. All data analysis was performed using the FieldTrip toolbox (Donders Institute for Brain, Cognition and Behaviour, Radboud University, the Netherlands. See http://fieldtriptoolbox.org) (Oostenveld, Fries, Maris, & Schoffelen, 2011) and custom MATLAB scripts.

### Task

Spatial memory was assessed using an “object location task” within a desktop virtual reality environment (Figure. 1A). Patients first navigated toward and memorized the location of four objects that sequentially appeared in the environment (‘encoding’). Patients were then cued with an image of one object (‘cue’), placed back in the environment and asked to navigate toward the remembered location of that object and make a button-press response (‘response’). The object then appeared in its correct location and the trial ended when they moved to the visible object (‘feedback’). Performance contrasts were computed by taking the median distance error across all trials for each participant and then comparing data between trials with error lower than the median (good trials) to those with error greater than the median (bad trials).

**Figure 1.**
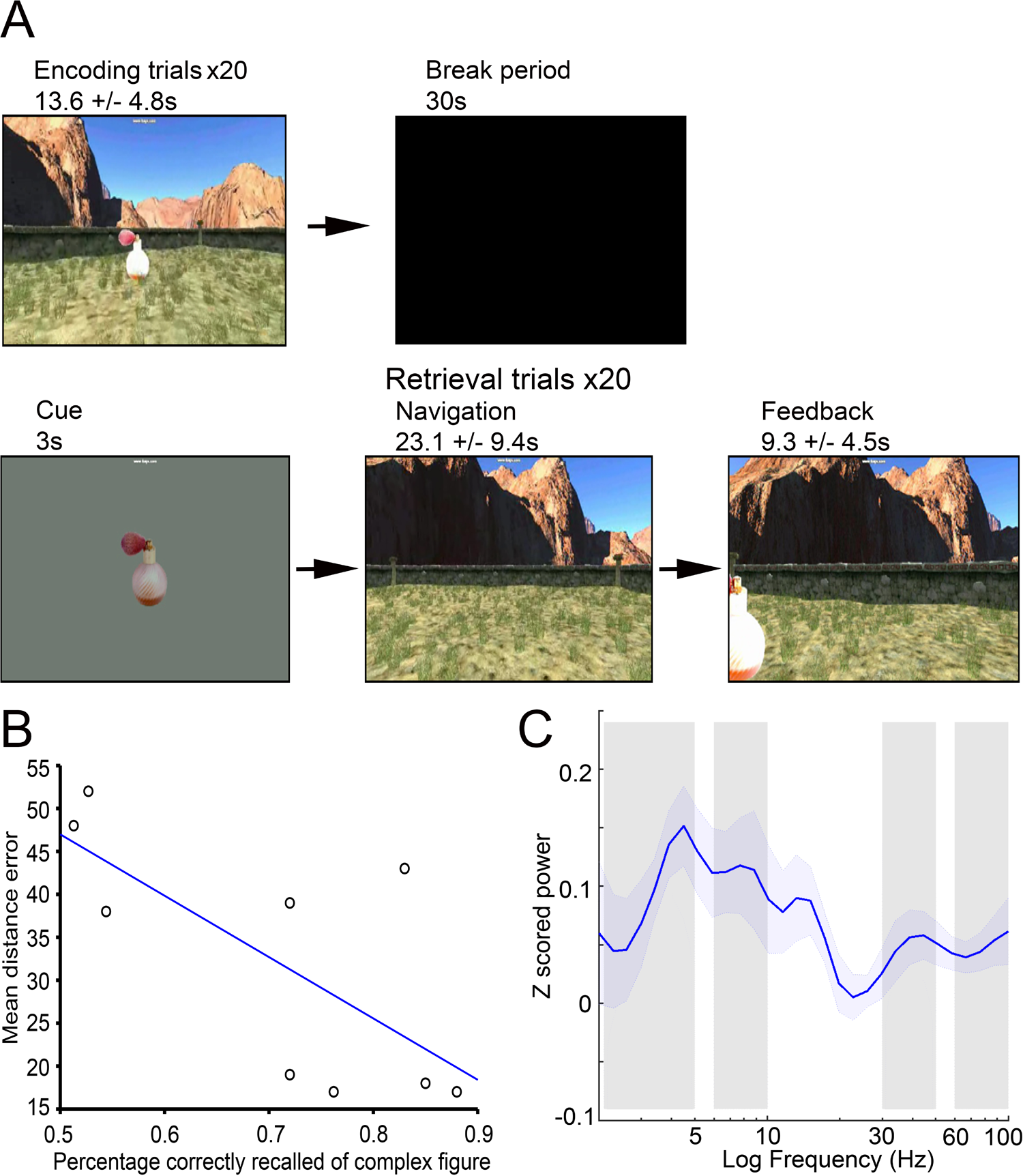
Increased theta and gamma power in the hippocampus during spatial memory retrieval. A. Schematic of the spatial memory task B. Median distance error in the spatial memory task versus assessment of complex figure recall. Line of best fit shown in blue C. Average z-scored power in the hippocampus during cue periods. Low (2–5 Hz) and high (6–9 Hz) theta bands and low (30-50Hz) and high (60-100Hz) gamma bands are marked in grey.

### Time–Frequency Analysis

Estimates of dynamic oscillatory power during periods of interest were obtained by convolving the EEG signal with a five-cycle Morlet wavelet. Time– frequency data were extracted from 2 s before the start of each cue period of interest to 2 s after the end of that period, and data from time windows before and after the period of interest were discarded after convolution to avoid edge effects. All trials that included interictal spikes or other artefacts, either within the period of interest or during the padding windows, were excluded from all analyses presented here. Each patient performed either one or two blocks, each consisting of 20 trials, providing a mean ± SD of 19.6 ± 11.9 trials for analysis, after artefact rejection. Power values were obtained for 30 logarithmically spaced frequency bands in the 2–100 Hz range. All power values from each contact were log transformed, and then z-scored using the mean and SD of log-transformed power values in each frequency band from artefact-free periods throughout the task.

To examine changes in oscillatory power within specific frequency bands and assess correlations between oscillatory power and task performance, dynamic estimates of log-transformed oscillatory power were averaged over the time and frequency windows of interest for each electrode contact. Mean power values were then averaged across all electrode contacts in the hippocampus to provide a single value for each patient. Changes in oscillatory power according to task demands were analysed using one sample t-tests, repeated-measures ANOVAs and post hoc one-sample t-tests with Bonferroni correction for multiple comparisons where appropriate.

### Phase amplitude coupling

PAC was tested for theta phase modulation of the amplitude of two specific gamma bands (30-50Hz and 60-100Hz; following Colgin et al., 2009). Phase-amplitude coupling was estimated using the phase-locking value, which is equal to the resultant vector length of the phase difference between low theta oscillations and the envelope (i.e. amplitude) of simultaneous gamma oscillations (Mormann et al., 2005). The resultant cross frequency coupling values were then z-scored across artefact-free periods from throughout the task, and comparisons made between good and bad trials. To establish whether changes in PAC across trials resulted purely from changes in the signal to noise ratio of low frequency power, we performed linear regression between low frequency power and PAC values across trials separately for each electrode contact. Beta coefficients were then averaged across all electrode contacts for each patient, allowing one-sample t-tests to be performed at the second level.

## Results

### 1. Object location task performance correlates with formal assessment of spatial memory

Prior to this study, all patients underwent formal neuropsychometry testing as pre-clinical evaluation for epilepsy surgery. As part of that testing, all patients (apart from one) were asked to perform a complex figure recall task from the BIRT Memory and Information Processing Battery (Coughlan, Oddy, & Crawford, 2007), as a measure of visuospatial memory, and a percentage accuracy score was given for performance. To establish the sensitivity of our object location task, we began by correlating patient performance during neuropsychometry with their mean distance error during that task. We found that there was a significant negative correlation between immediate recall of the complex figure and distance error in our task (Pearson’s r = -0.71, p=0.032; Figure 1B). This suggests that our object location task was sensitive in discerning the spatial memory abilities of our subjects.

### 2. Increased theta and gamma power in hippocampus during retrieval

Next, we examined low frequency oscillatory power on hippocampal contacts during the 3s cue period, when participants were asked to retrieve the location of an object prior to being replaced in the virtual environment. We found that average z-scored theta power in both the low (2-5Hz; t(9)=4.36, p=0.0024) and high (6-9Hz; t(9)=3.04, p=0.016) frequency bands were significantly higher during the cue period. This suggests that low and high theta oscillations in the hippocampus are engaged as a result of spatial memory retrieval. Next, we focussed on high frequency oscillatory power in the 30-50Hz and 60-100Hz gamma bands. We found that average z-scored gamma power in both the low (30-50Hz; t(9)=2.34, p=0.047) and high (60-100Hz, t(9)=2.55, p=0.034) frequency bands were significantly higher during the cue period (Figure 1C). This suggests that hippocampal gamma oscillations are also engaged by spatial memory retrieval.

### 3. Increased theta power in hippocampus during retrieval predicts performance

Next, in order to establish whether increased theta and / or gamma power in the hippocampus were associated with task performance, we conducted a repeated measures ANOVA with factors of performance (good v bad trials) and theta frequency band (low v high). We found a significant effect of performance (F(1,35)=6.4, p=0.035), but no interaction (p=0.68), driven by increased low and high theta power during good trials (Figure 2A, B). Repeating this analysis with factors of performance (good v bad) and gamma frequency band (low v high) revealed no significant effect of performance or interaction (all p>0.094). This suggests that increased theta power within the hippocampus is associated with accurate performance of the object location task, while gamma power does not vary between good and bad trials.

**Figure 2.**
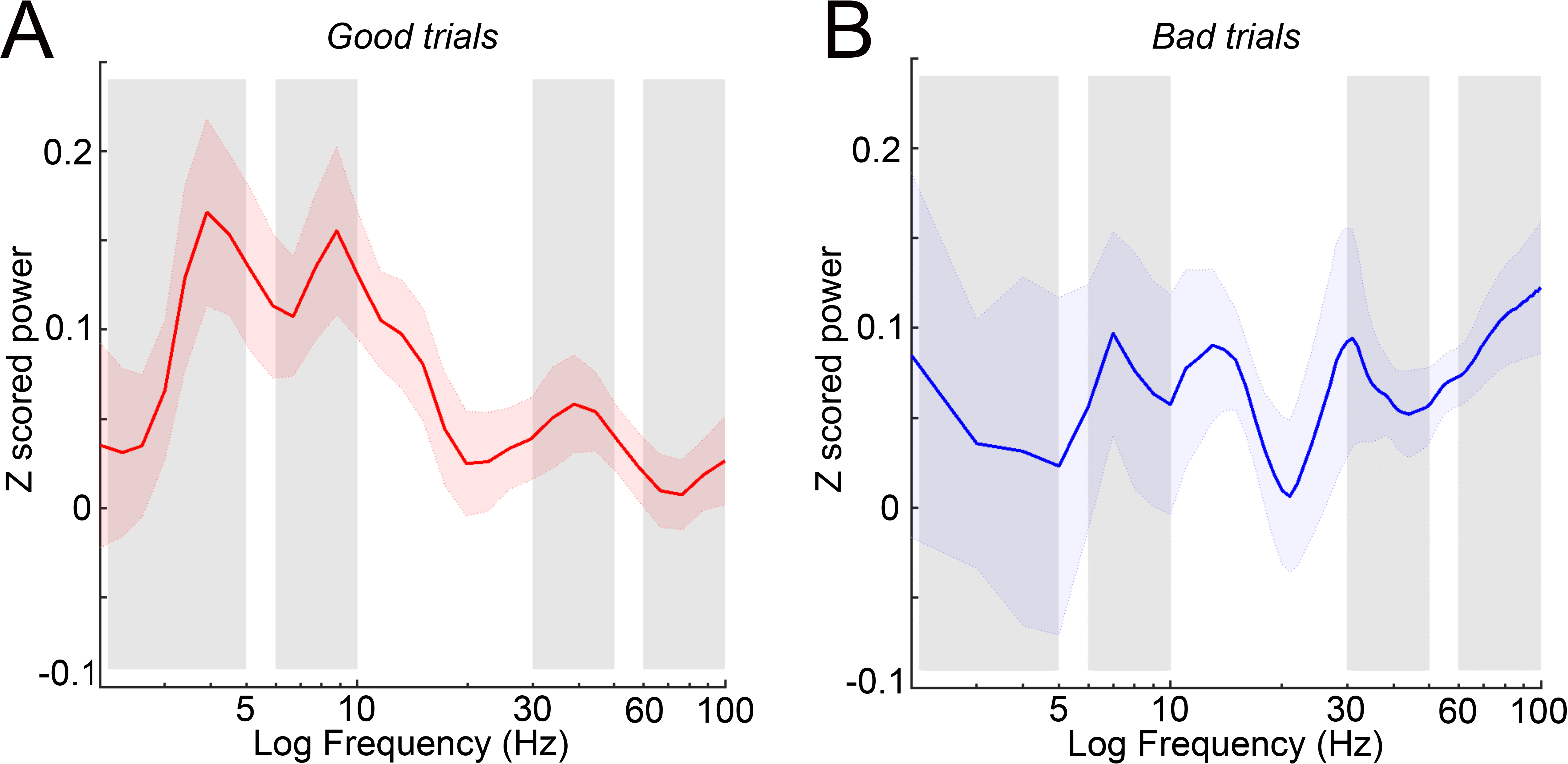
Low and high theta power in the hippocampus are associated with improved performance. A. Average z-scored power in the hippocampus during cue periods of good trials, with low (2–5 Hz) and high (6–9 Hz) theta bands and low (30-50Hz) and high (60-100Hz) gamma bands marked in grey. B. Average z-scored power in the hippocampus during cue periods of bad trials.

### 4. Increased theta-gamma phase amplitude coupling in hippocampus during retrieval predicts performance

To further dissect the role of low and high theta oscillations in spatial memory retrieval, we next examined changes in phase-amplitude coupling (PAC) between low or high frequency theta phase and low or high frequency gamma amplitude. First, we conducted a repeated measures ANOVA for all trials with factors of theta band (2-5Hz v 6-9Hz) and gamma band (30-50Hz v 60-100Hz), but found no significant differences in z-scored PAC between any pair of frequency bands (all p>0.36). In addition, z-scored PAC values between any pair of theta and gamma frequency bands were not significantly different from zero (all p>0.15), suggesting that there was no overall change in theta-gamma PAC during spatial memory retrieval.

Next, to examine if PAC in the hippocampus was relevant for task performance we performed a three way ANOVA with factors of performance (good v bad), theta band (2-5Hz v 6-9Hz) and gamma band (30-50Hz v 60-100Hz). We found that PAC differed significantly between good and bad trials (F(1,63)=9.21, p=0.005), with a significant interaction between good/bad trials and low/high theta (F(1,63)=4.88, p=0.031), but no other interactions (p>0.17). Subsequent analysis indicated that these results were driven by the increased modulation of both low and high gamma amplitude by the phase of low theta band oscillations during good trials (t(9)=2.2 p=0.02; Figure 3A-D). In order to ascertain whether this increase in PAC was solely a reflection of increased low theta power, and therefore signal-to-noise ratio, we examined whether low theta power and PAC values correlated across trials on each electrode contact, but found no evidence for a significant linear relationship (p=0.23). This suggests that increased low theta phase modulation of both low and high gamma amplitude in the hippocampus is associated with improved task performance, independent of concurrent changes in low theta power.

**Figure 3.**
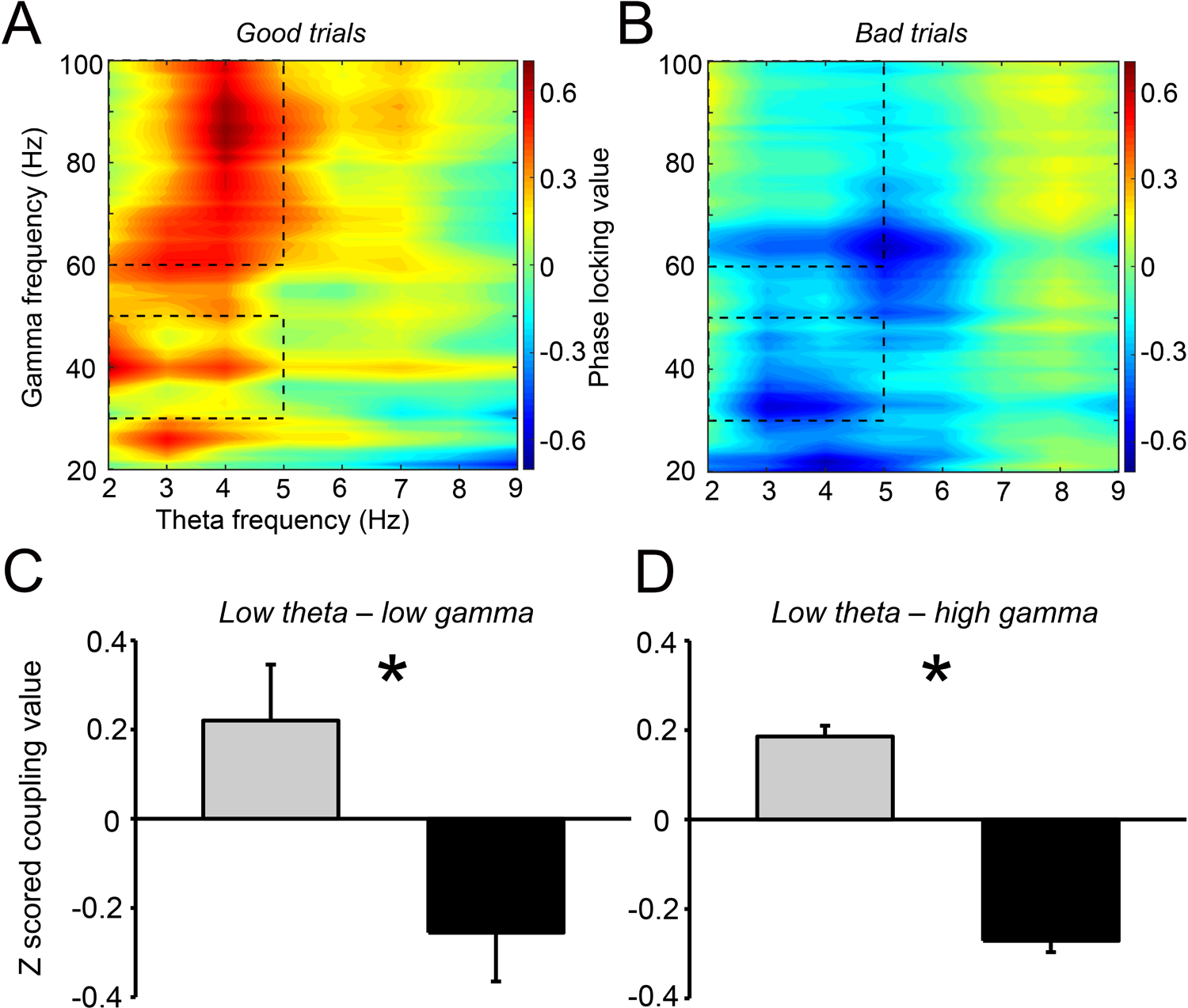
Increased low theta phase modulation of gamma amplitude is associated with improved performance. A. Cross frequency spectrogram for good trials; boxed regions highlight significant coupling between 2-5Hz low theta phase and 30-50Hz low and 60-100Hz high gamma amplitude B. Cross frequency spectrogram for bad trials C. Average z-scored PAC between 2-5Hz low theta phase and 30-50Hz low gamma amplitude within the hippocampus in good (grey) and bad (black) trials. D. Average z-scored PAC between 2-5Hz low theta phase and 60-100Hz high gamma amplitude within the hippocampus in good (grey) and bad (black) trials

## Discussion

Although separate low (2-5Hz) and high (6-9Hz) theta bands have been observed during mnemonic function in humans, their specific function has remained unclear. Here, we have demonstrated distinct roles for low and high theta within the hippocampus during performance of a spatial memory task. It appears that during spatial memory retrieval (i.e. during cue periods), both low and high theta power and low and high gamma power are increased in the hippocampus. We had previously shown that hippocampal theta power was indicative of movement onset in this task (Bush et al., 2017). Theta power is also associated with task performance, as previously reported in MEG (Kaplan et al., 2012) and intracranial EEG (Miller et al., 2018) studies. Moreover, we found that increased modulation of low (30-50Hz) and high gamma (60-100Hz) amplitude by low theta phase in the hippocampus is associated with improved task performance. Indeed, rodent models suggest that these gamma frequency bands mediate the routing and temporal segregation of inputs to the hippocampal CA1 region from different sources (Colgin et al., 2009). In addition, human studies have demonstrated a relationship between hippocampal subfield gamma power and spatial memory precision in a different object-location task (Stevenson et al., 2018).

In summary theta band oscillations are functionally relevant to spatial memory retrieval, while low theta-gamma phase amplitude coupling has a role in accurate spatial memory performance.

## Acknowledgements

This work was supported by the Department of Health’s National Institute for Health Research, UCL/UCL Biomedical Research Centre, Wellcome Trust, UK Medical Research Council, European Research Council, Epilepsy Research UK, and Academy of Medical Sciences.

We thank all patients who participated in this study.

## Figure legends

Table 1. Demographics and Epilepsy History of Patients for Spatial Memory Task. F=female, M=male, R=right, L=left, B=bilateral

